# Complex changes in serum protein levels in COVID-19 convalescents

**DOI:** 10.1101/2022.10.26.513886

**Authors:** Smruti Pushalkar, Shaohuan Wu, Shuvadeep Maity, Matthew Pressler, Justin Rendleman, Burcu Vitrinel, Lauren Jeffrey, Ryah Abdelhadi, Mechi Chen, Ted Ross, Michael Carlock, Hyungwon Choi, Christine Vogel

## Abstract

The COVID-19 pandemic, triggered by severe acute respiratory syndrome coronavirus 2, has affected millions of people worldwide. Much research has been dedicated to our understanding of COVID-19 disease heterogeneity and severity, but less is known about recovery associated changes. To address this gap in knowledge, we quantified the proteome from serum samples from 29 COVID-19 convalescents and 29 age-, race-, and sex-matched healthy controls. Samples were acquired within the first months of the pandemic. Many proteins from pathways known to change during acute COVID-19 illness, such as from the complement cascade, coagulation system, inflammation and adaptive immune system, had returned to levels seen in healthy controls. In comparison, we identified 22 and 15 proteins with significantly elevated and lowered levels, respectively, amongst COVID-19 convalescents compared to healthy controls. Some of the changes were similar to those observed for the acute phase of the disease, i.e. elevated levels of proteins from hemolysis, the adaptive immune systems, and inflammation. In contrast, some alterations opposed those in the acute phase, e.g. elevated levels of CETP and APOA1 which function in lipid/cholesterol metabolism, and decreased levels of proteins from the complement cascade (e.g. C1R, C1S, and VWF), the coagulation system (e.g. THBS1 and VWF), and the regulation of the actin cytoskeleton (e.g. PFN1 and CFL1) amongst COVID-19 convalescents. We speculate that some of these shifts might originate from a transient decrease in platelet counts upon recovery from the disease. Finally, we observed race-specific changes, e.g. with respect to immunoglobulins and proteins related to cholesterol metabolism.

## Introduction

The COVID-19 pandemic has affected more than 607 million people worldwide with approximately 6.5 million deaths (World Health Organization). Caused by severe acute respiratory syndrome coronavirus 2 (SARS-CoV-2), the disease is highly infective and exhibits an extensive clinical heterogeneity, from asymptomatic to symptomatic disease states ^1^. While the primary manifestation of COVID-19 is in the respiratory tract, there is an increased risk of other life-threatening pathologies such as pulmonary embolism, myocardial infarction, and ischemic stroke with frequent venous and arterial thromboembolisms with the severity of disease ^2^. Similarly, recovery from COVID-19 has displayed enormous heterogeneity, ranging from disappearance of symptoms within a few days to establishment of ‘long COVID’, also known as ‘post-acute sequelae of SARS-CoV-2’ (PASC) marked by a broad spectrum of the ongoing physiological changes ^3,4^.

Much work has been done to characterize the molecular changes during the acute phase of the disease, e.g. in patient plasma and serum samples ^5–8^. Both untargeted and targeted proteomics approaches have identified dysregulation of various pathways including lipid homeostasis, immunoglobulins, inflammatory and antiviral cytokines, chemokines of innate and adaptive immunity, as well as complement and coagulation cascades ^4,5,9–11^. These studies also observed platelet degranulation, lymphocyte apoptosis in serum, likely due to the viral mode of entry into the cells ^4,5,9,10^. During acute infection, overproduction of proinflammatory cytokines (IL-6, IL-1β, and TNF-α) induces a ‘cytokine storm’ elevating the risk of clot formation, platelet activation, and ultimately hypoxia and multiorgan failure leading to high patient mortality ^9,12^. Accordingly, serum proteomics using a highly sensitive targeted assay identified proteins of inflammation, cardiometabolic, and neurologic diseases to contribute to disease severity ^13^. Other studies found an expansion in regulatory proteins of coagulation (APOH, FN1, HRG, KNG1, and PLG) and lipid homeostasis (APOA1, APOC1, APOC2, APOC3, and PON1) in serum as the disease progressed ^5^. COVID-19 plasma samples also demonstrated extensive changes in several key protein modifications, such as glycosylation, phosphorylation, citrullination and arginylation during the acute phase of the disease ^14^. Even serum from COVID-19 infected asymptomatic individuals showed altered levels of coagulation and inflammation, such as fibrinogen, von willebrand factor (VWF), and thrombospondin-1 (TSP1)^3^. In comparison, long COVID/PASC patients appear to have altered levels of autoantibodies, localized inflammation, and reactivation of latent pathogens ^4^. In particular, patients of neuro-PASC exhibit plasma proteomes highly distinct from COVID-19 convalescents who have no lingering symptoms, e.g. presenting markers of DNA repair, oxidative stress, and neutrophil degranulation ^15^.

In comparison, information on COVID-19 convalescence is much less available, in particular from patients without vaccination or prior SARS-CoV-2 infections. One study examining blood samples from severe COVID-19 patients upon recovery observed elevated erythrocyte sedimentation rates, increased levels of C-reactive protein (CRP), and reduced levels of serum albumin ^16^. Another study showed that carbonic anhydrase 1 (CA1) was still bound to immunoglobulin IgA in COVID-19 patients within 2 weeks of recovery, unlike in any healthy vaccinated or unvaccinated healthy subjects, or in COVID-19 patients after 6 months of recovery ^17^. Another study identified lipid, atherosclerosis and cholesterol metabolism pathways, complement and coagulation cascades, autophagy, and lysosomal function for at least six weeks upon recovery from infection with the ancestral SARS-CoV-2 virus ^18^. Similar substantial changes in the plasma proteome were found in hospitalized COVID-19 patients even six months after discharge ^19^.

To understand the broader impact of COVID-19 after recovery, we profiled the proteome in 29 serum samples from a unique collection of COVID-19 convalescents and 29 samples from age-, sex-, and race-matched healthy controls. All COVID-19 convalescents had tested positive for SARS-CoV2 in March or April 2020, i.e. during the first months of the pandemic. All convalescents were symptomless at the time of sample collection, i.e. considered outside the acute phase of the disease. We used mass spectrometry to quantify protein levels in the soluble fraction of the blood serum in an untargeted fashion and used linear regression models to deconvolute the effects of demographic parameters in differentially abundant serum proteins. We identified pathways whose member proteins had returned to the pre-infection levels, and pathways with member proteins that were still altered either consistent with or opposite to changes observed during the acute infection.

## Results

### Quantitation of 334 serum proteins

We quantified proteins from serum samples of 29 COVID-19 convalescents and age-, sex- and race-matched 29 healthy individuals. **Figure 1A** shows the experimental outline; **Supplementary Table S1** describes the cohort demographics, including known information on the acute phase of COVID-19. Participant age ranged from 22 to 61 years, with a median (interquartile range, IQR) of 44 (20) years. The male to female ratio was 1.2. About 55% of the individuals reported to have had symptoms at the time of the acute COVID-19 infection while the remaining individuals reported no symptoms.

**Figure 1.**
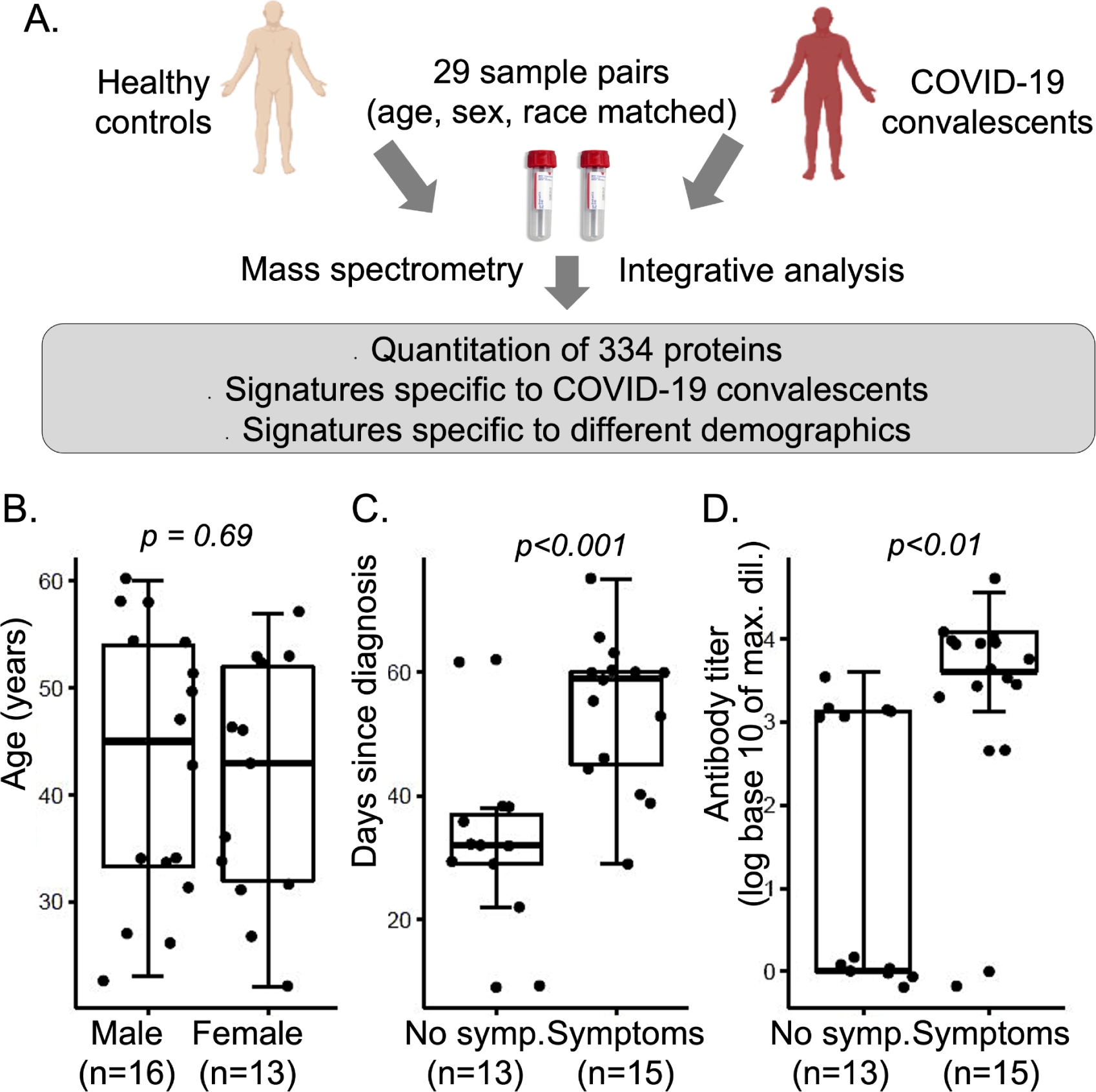
Experimental design. **A.** Overview of experimental design providing proteomic analysis of serum samples from COVID-19 convalescents with age, sex, and race matched healthy controls. **B. to D.** Selection of demographic factors describing the 29 COVID-19 convalescents: **B.** Distribution of age between male and female cases. **C.** and **D.** Distribution of Days since diagnosis and Antibody titer levels respectively, split by self-reported presence of symptoms during the acute phase. All demographic data are provided in **Supplementary Data File S1** and summarized in **Supplementary Table S1**. P-values were derived from t-tests. Symp. - symptoms.

The serum samples from COVID-19 convalescents were collected 9 to 70 days after diagnosis (Days since diagnosis). Individuals had been diagnosed by PCR test in March/April 2020, i.e. a few months after the beginning of the pandemic. Participants were recruited based on opportunity without further exclusion/inclusion criteria, were unvaccinated and experienced their first infection with SARS-CoV-2. None of the individuals had been hospitalized or had lingering symptoms at the time of sample collection. Thirteen individuals (45%) reported that they experienced no symptoms during the acute phase of the infection. Fifteen individuals (52%) reported symptoms during the acute infection, with fever, shortness of breath, loss of taste, and coughing amongst the most frequent (**Supplementary Table S1**). Due to the small cohort size and range of symptoms, we considered COVID-19 convalescents only as symptomatic or asymptomatic without further distinction.

Serum antibody titers were measured at the time of sample collection. About 24% of the patients showed no antibody response whereas 3%, 21%, 21% and 24% patients displayed low, moderate, high and very high antibody response respectively, as defined in **Supplementary Table S1**.

Figures 1B-D describe relationships between selected demographics. The age distributions between male and female participants were similar **(Figure 1B)**. Individuals who reported symptoms during the acute phase of the infection had a significantly longer period until sample collection (p-value < 0.05, Figure 1C): the median (IQR) Days since diagnosis were 59 (15) for symptomatic and 32 (8) for asymptomatic individuals (**Supplementary Table S1**). Similarly, individuals with symptoms had significantly higher antibody titers (Figure 1D) than individuals without symptoms (p-value < 0.05), and Days since diagnosis and antibody titers correlated substantially (R2=0.44, **Supplementary Figure S1A**). One possible explanation for these correlations was that participants entered the study at a time based on severity of the infection (i.e. the extent of symptoms) and length of recuperation time: more severely ill individuals provided serum samples at a later time than less ill individuals. Conversely, more severely ill individuals might have produced higher antibody titers, even if collected at a later point after recovery. Based on the convoluted relationships between Days since diagnosis, Symptoms, and Titer levels, the wide range in values, the lack of additional information, and the small cohort size, we refrained from extensive analysis of these characteristics except for what is described below.

We quantified a total of 334 proteins across all samples as depicted in Figure 2. About half of the proteins fall into three functional categories: immunoglobulins, complement cascade and high-density lipoproteins (32%, 9%, and 5%, respectively). We grouped the proteins into 20 clusters based on their levels across the samples and the similarity in expression patterns (see **Methods**). The cluster number was chosen based on the total number of proteins. Twelve clusters contained one or more proteins with statistically significant differences between COVID-19 convalescents and healthy controls; and we focus discussion on these 12 clusters below. We tested the 12 clusters for function enrichment and show example proteins in figures.

**Figure 2.**
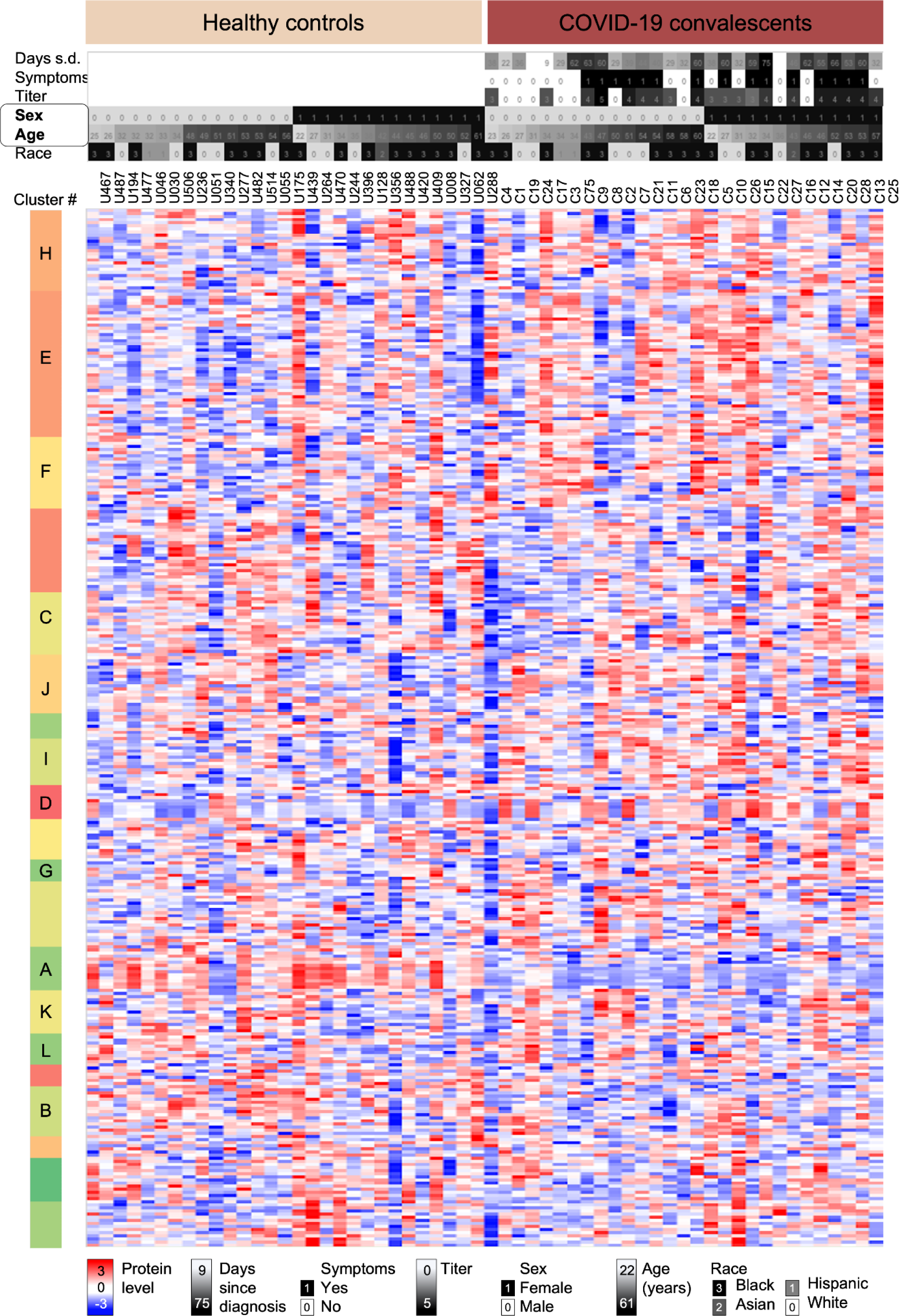
Normalized levels of 334 serum proteins. Heatmap depicts normalized log_10_-transformed levels for 334 proteins in sera from healthy controls and COVID-19 convalescents sorted according to sex and age. The upper panel provides additional sample information such as days passed since diagnosis of an acute SARS-CoV-2 infection and sample collection, self-reported presence of symptoms during the acute phase, antibody titer levels (log_10_-transformed), sex, age, and race of the individual. All demographic data are provided in **Supplementary Data File S1** and summarized in **Supplementary Table S1**. After hierarchical clustering we split the data into 20 clusters with specific protein expression patterns; letters indicate clusters discussed in detail in the text. Each cluster was examined for trends in expression differences and function enrichment (see **Methods**). S.d. - since diagnosis.

All protein measurements underwent extensive normalization (see **Methods**). After normalization, we evaluated the quality of the normalized measurements through different methods. First, we analyzed the coefficient of variance (CoV) for the four technical matches for all normalized protein measurements across the QC samples (**Supplementary Figure S1B**). The median CoV was <30% in all four batches, which is consistent with what is expected from untargeted proteomics analyses. Only 40 proteins showed an average CoV>50% in the QC samples (**Supplementary Data File S1**); their quantification in the cohort samples is less reliable, and the proteins are marked in all figures with a *. Further, we examined the sample distribution across the first and second principal component, which showed successful clustering of all QC samples (**Supplementary Figure S1C**). Finally, as an additional quality control, we examined several example proteins for their levels amongst demographic subsets (**Supplementary Figures S2** and **S3**). The proteins showed the expected sex and body weight related differences.

### Altered levels of components of the innate and adaptive immune system

First, we examined the overall difference in protein levels between samples from healthy controls and COVID-19 convalescents, regardless of the individuals’ demographics. We conducted partial least squares discriminant analysis with the protein levels (Figure 3A). The major components explained 19% of the variability and separated the data into two distinct clusters comprising the two cohorts. This separation confirmed that the proteomics data captured differences between COVID-19 convalescents and healthy controls.

**Figure 3.**
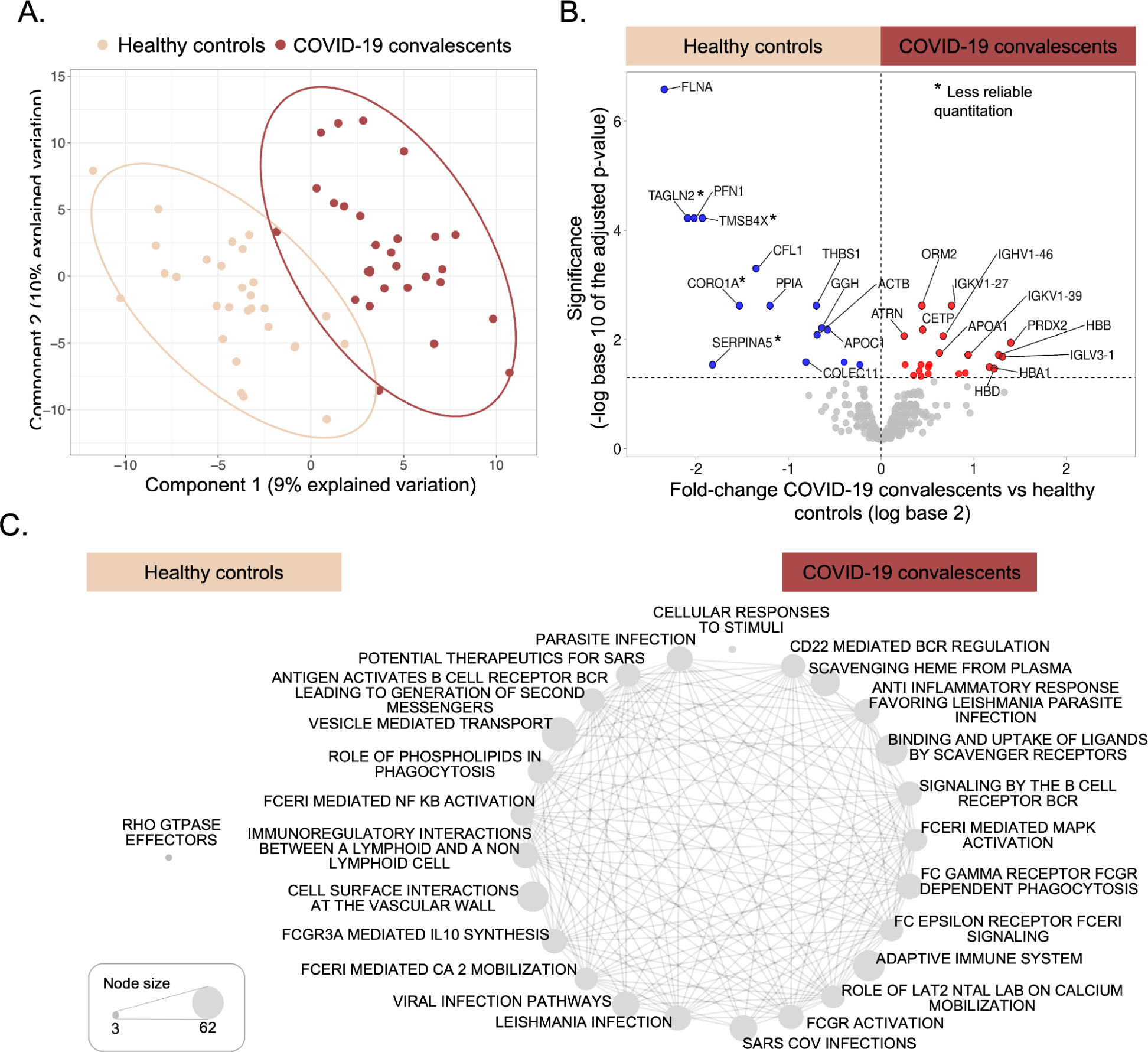
Differential protein levels in COVID-19 convalescents and healthy controls. **A**. Partial least square discriminant analysis depicts segregation between the two cohorts. **Supplementary Figure S1** shows further characteristics of quality control: the distributions of the Coefficients of Variance (**S1B**) and the first and second principal component including Quality Control samples and sample labels in a plot analogous to this one (**S2C**). **B.** The volcano plot indicates fold changes and corresponding adjusted p-values of protein levels between COVID-19 convalescents and healthy controls. Colored dots represent proteins with significantly higher levels amongst the COVID-19 convalescents (red) or healthy controls (blue), respectively (adjusted p-value <= 0.05). The expression patterns of these proteins are also listed in **Supplementary Figure S4**. A * marks proteins with less reliable quantitation as determined by a high Coefficient of Variance across quality control samples (>50%). **C**. The networks visualize the function enrichment amongst proteins ranked by their directed adjusted p-value (if the observed log_10_-transformed fold change was positive, we calculated 1-p; if negative, we calculated -(1-p)). Healthy control samples have only one enriched function (false discovery rate < 0.05). Details of network construction are provided in the **Methods** section.

The volcano plot in Figure 3B depicts the results from the overall comparison of protein levels between the two sample sets. Dots with red and blue colors in Figure 3B marked the 22 and 15 proteins that were significantly elevated or decreased in the serum from COVID-19 convalescents and healthy controls, respectively (adjusted p-value < 0.05). **Supplementary Figure S4** shows the levels of these proteins measured in each sample. The COVID-19 convalescents had significantly elevated immunoglobulins, Orosomucoid 2 (ORM2), peroxiredoxin-2 (PRDX2), hemoglobin subunits (HBD, HBB and HBA1), as well as proteins involved in cholesterol transport such as cholesteryl ester transfer protein (CETP) and apolipoprotein A1 (APOA1). In comparison, healthy controls had significantly elevated levels of actin cytoskeleton signaling proteins, e.g. filamin (FLNA), profilin (PFN1), cofilin (CFL1), and actin beta (ACTB).

We rank-ordered proteins according to their differential levels amongst COVID-19 convalescents vs. controls and analyzed them for enriched protein functions (see **Methods**). Significantly enriched functions and pathways are represented in Figure 3C (false discovery rate < 0.05). Proteins enriched healthy controls are biased with respect to only one pathway with three protein members (single node in Figure 3C). In comparison, proteins with higher levels in COVID-19 convalescents are enriched in a large number of pathways with many shared protein members (edges in Figure 3C). The pathways relate to (viral) infection, the innate and adaptive immune response, signaling cascades, inflammation, and general cellular processes. Due to the large number of overlapping categories, we subsetted the analysis below into proteins with differential expression patterns *in addition* to those observed between COVID-19 convalescents and healthy controls.

The volcano plot in Figure 3B also shows the proteins with similar levels in both healthy controls and COVID-19 convalescents (gray dots below the 0.05 threshold). These proteins either did not change during the acute infection or had returned to normal levels at the time of analysis. **Table 1A** shows the subset of the proteins with similar levels in both healthy controls and COVID-19 convalescents, but altered levels during acute COVID-19, as known from literature. The proteins include members of the complement (e.g. C2, C3, C4A) and coagulation (e.g. F5, F10) cascade, the adaptive immune system, metabolism (e.g. LPA, PON1) and inflammation (e.g. ORM1, S100A8, and S100A9). **Supplementary Figure S5** shows the expression patterns for additional proteins from the complement system, coagulation cascade, and from inflammation. As both members of the complement system and coagulation cascade are heavily dysregulated during the acute phase of the infection ^20–22^. An analysis of COVID-19 patients during the acute infections up to six weeks of recovery also revealed elevated levels of proteins of the complement system and coagulation cascade ^18^. It is possible that in our study they had changed in the COVID-19 convalescents during the acute phase but had returned to levels similar to those observed in healthy individuals by the time of analysis.

**Table 1.** Summary of the results in relationship to changes reported in literature. The table lists example pathways and proteins whose levels were altered during the acute phase of COVID-19 as known from literature and the levels as measured in the group of COVID-19 convalescents in this study. Example proteins had been chosen based on their citation in literature (see text for references) and their membership in the respective functions/pathways. **A.** Protein levels in COVID-19 convalescents are similar to those measured in healthy individuals in this study. **B.** Protein levels in COVID-19 convalescents are similar to those observed in acute COVID-19 cases. **C.** Protein levels in COVID-19 convalescents differ from those observed in acute COVID-19 cases and from those measured in healthy individuals in this study.

| Pathway | Example proteins | Known: | This Study: |
| --- | --- | --- | --- |
|  |  | Acute phase<br>COVID-19 | COVID-19<br>convalescents |
| A. Levels in COVID-19 convalescents similar to those in healthy individuals |  |  |  |
| Complement cascade | C2, C3, C4A, C4B, C8,<br>C9, MASP1, MBL2, FCN2 | Elevated | Healthy |
| Coagulation system/thrombosis | F5, F10, FGB, FGG, KNG1 | Elevated | Healthy |
|  | F13B | Lower |  |
| Adaptive immune system | IGLV3-25, IGKV1-5 | Elevated | Healthy |
| Metabolism | LPA, PON1 | Elevated | Healthy |
|  | APOA2, APOA4 | Lower |  |
| Inflammation/Immune response | CRP, LDH, ORM1, S100A8, S100A9 | Elevated | Healthy |
| **B. Levels in COVID-19 convalescents different from those in healthy individuals but similar to those in acute COVID-19 patients** |  |  |  |
| Hemolysis | HBA1, HBB, HBD, PRDX2, CA1 | Elevated | Elevated |
| Adaptive immune system | IGHG1, IGHV1-24, IGHV1-46, IGHV1-69, IGKV3D-15, IGLV3-19 | Elevated | Elevated |
| Inflammation/Immune response | ORM2, SAA4 | Elevated | Elevated |
| **C. Levels in COVID-19 convalescents different from those in healthy individuals and from those in acute COVID-19 patients** |  |  |  |
| Complement cascade | C1R, C1S, CFI, COLEC11, VWF | Elevated | Lower |
| Coagulation system/thrombosis | VWF, FN1, THBS1 | Elevated | Lower |
| Actin cytoskeleton/Cell adhesion/Platelet degranulation | PFN1, CFL1, FLNA, VWF | Elevated | Lower |
| Metabolism | CETP, APOA1 | Lower | Elevated |

We also tested for differences in protein modifications between the two groups. While we did not enrich proteins with post-translational modifications, the type of mass spectrometry data we collected allowed for a retrospect analysis for modified peptides. To do so, we constructed computational libraries scanning all data for the occurrence of several frequently occurring modifications, i.e. mono- and di-hexose, phosphorylation, and mono- and di-methylation, and then used the library to analyze the cohort samples. We observed hexose addition (glycosylation) most frequently (**Supplementary Data File S2**): three of 55 hexose-modified peptides were significantly more abundant in COVID-19 convalescents than in healthy controls, when normalized for the levels of the respective unmodified peptides (adjusted p-value < 0.05; **Supplementary Figure S6**). The three peptides originated from albumin (ALB) and immunoglobulin heavy constant alpha 1 (IGHA1); these two proteins did not show significantly differential levels across the two cohorts. Our results were consistent with extensive serum glycosylation observed in COVID-19 patients ^23,24^. Other modifications affected only a few peptides in our data.

### Impact of demographic factors on serum protein signatures

Next, we tested for the impact of demographics, i.e. Days since diagnosis, Symptoms, Age, Sex, and Race of the individuals on the serum protein levels on the respective COVID-19 states to extract patterns beyond the simple difference between COVID-19 convalescents and healthy individuals. To do so, we used a variety of models (see **Methods**). Due to the correlation between Titer and Symptoms as well as Days since diagnosis, we excluded Titer levels from the analyses. We tested i) protein levels in healthy individuals; ii) protein levels in COVID-19 convalescents; and iii) log base 2 ratios of the protein levels in the COVID-19 cases versus the matched healthy controls. Figures 4 and **5** show the results, with proteins grouped according to the clusters identified in Figure 2. Example proteins for each cluster were chosen based on their membership in the functions in the respective cluster. Results from all analyses are shown in **Supplementary Data File S1**.

**Figure 4.**
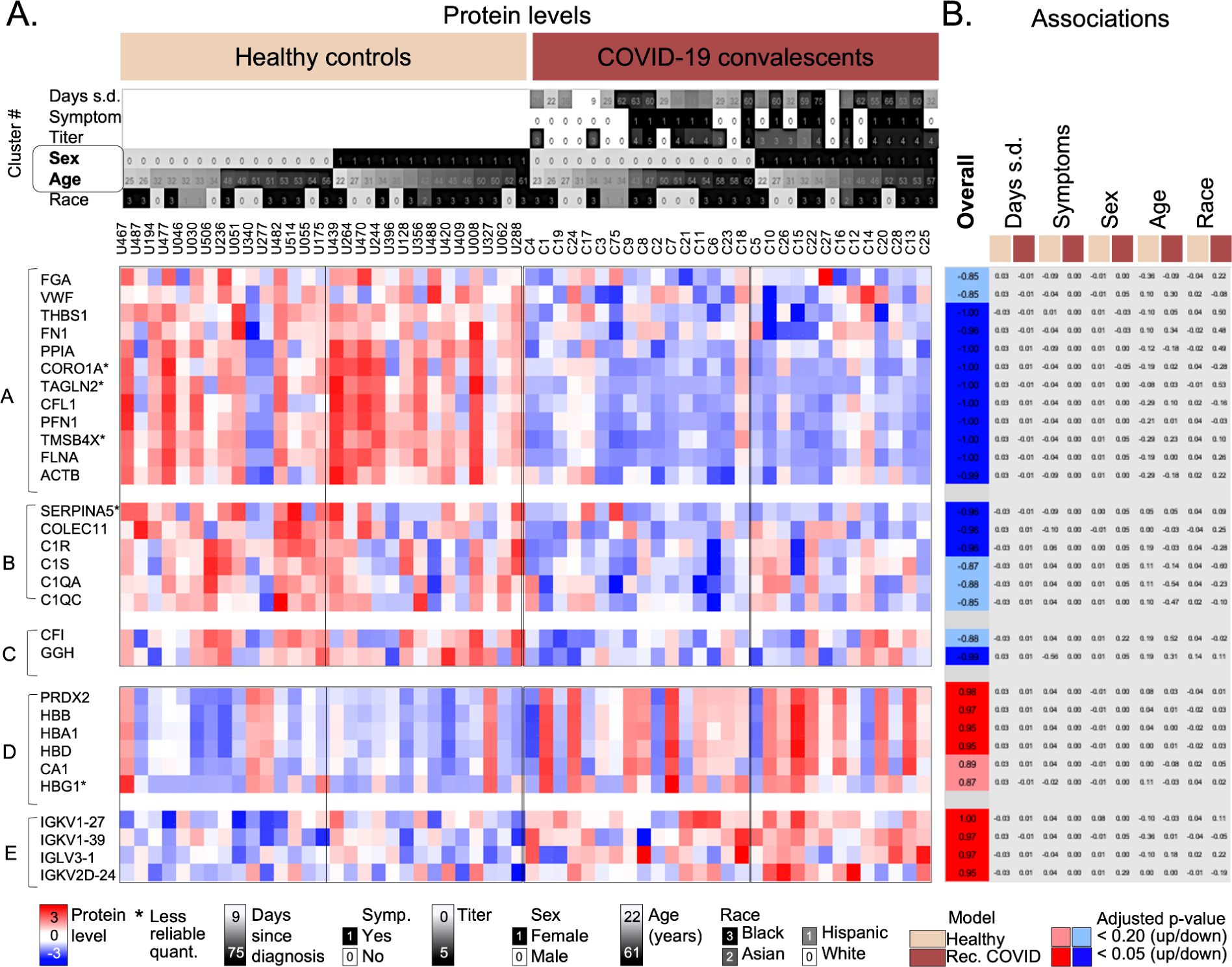
Levels of proteins with differences between healthy controls and COVID-19 convalescents independent of demographic factors. Heatmap depicting example proteins from five clusters (marked in **Figure 2**). The clusters were selected based on the significant differences between COVID-19 convalescents and healthy controls. Each cluster was analyzed for enriched functions, and examples were selected based on these functions (see **Methods**). Function enrichments were as follows: A: Cell adhesion, platelet degranulation; B and C: Innate immunity, complement system; D: Hemoglobin, adaptive immunity, immunoglobulins; and E: Adaptive immunity, immunoglobulins, complement system. Example proteins were selected based on the statistical significance of the overall difference between protein levels in COVID-19 convalescents and healthy controls (with the adjusted p-value < 0.20 or < 0.05) and based on their relevance for the observed functional enrichment in each cluster. Panel **A**. shows the color-coded protein levels with samples sorted according to sex and age. Panel **B**. shows the significance values for two different models. Significance values (adjusted p-values) were transformed as follows: if the observed log_10_-transformed fold change was positive, we calculated 1-p; if negative, we calculated -(1-p). Dark colors indicate adjusted p-value <0.05; light colors adjusted p-value < 0.20; gray: no significance. Columns represent comparisons with significant differences: the Overall difference between COVID-19 convalescents and healthy controls; and the impact of Days since diagnosis, presence of Symptoms, Sex, Age, and Race in a multivariate model using the healthy controls (beige) or COVID-19 convalescents (brown), respectively. The demographic data, protein levels, and results of all models are provided in **Supplementary Data File S1**. D.s.d. - Days since diagnosis; Titer - log_10-_transformed antibody titer as defined in methods; * Less reliable quantitation as determined by a high coefficient of variance across quality control samples (>50%)

**Figure 5.**
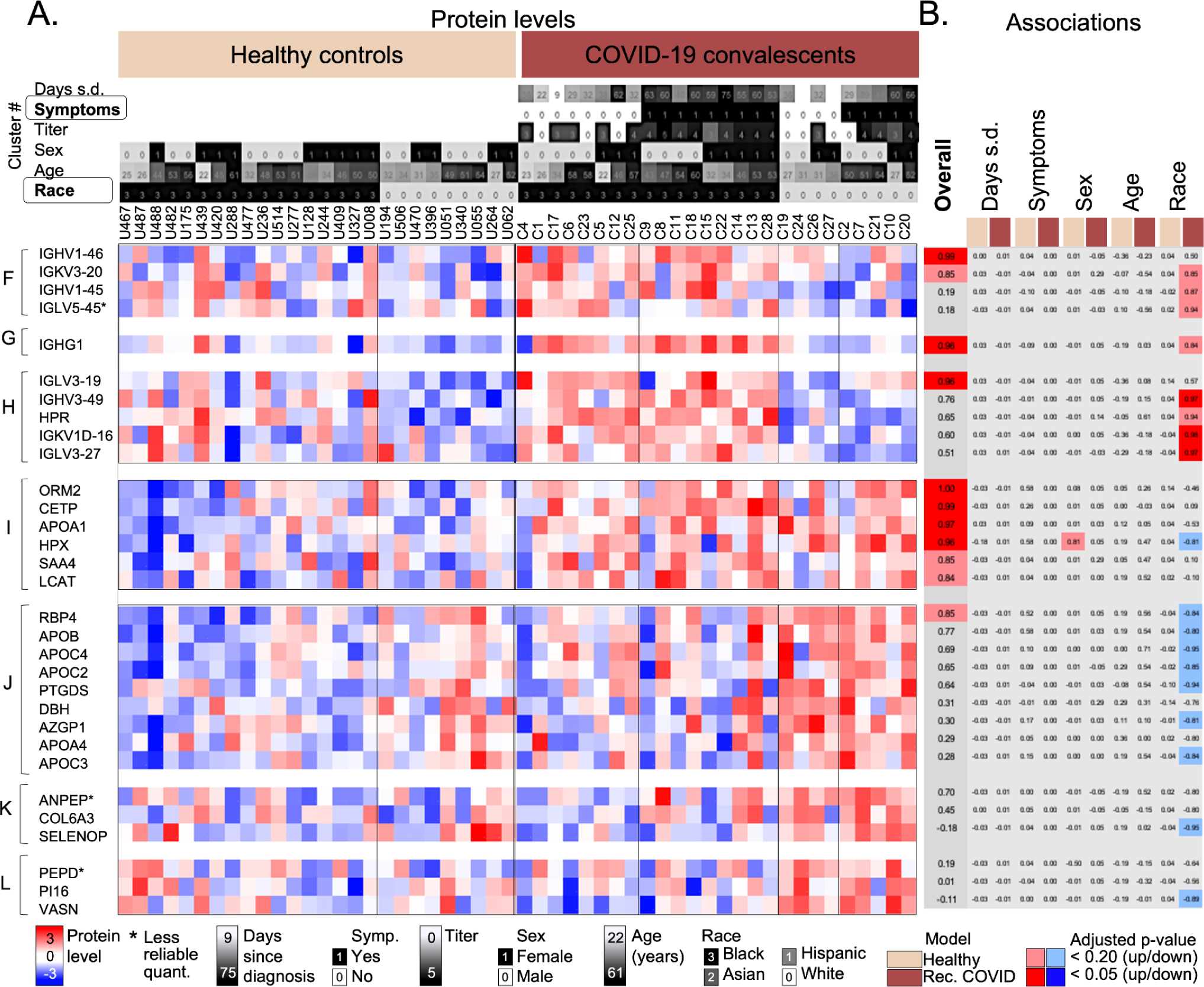
Levels of proteins in COVID-19 convalescents and healthy controls with interactions with other factors. Heatmap depicting example proteins from five clusters (marked in **Figure 2**). The clusters were selected based on the significant differences between COVID-19 convalescents and healthy controls. Each cluster was analyzed for enriched functions, and examples were selected based on these functions (see **Methods**). Function enrichments were as follows: F: Adaptive immunity, immunoglobulins; H: Adaptive immunity, immunoglobulins; I: Cholesterol transport; J: Lipid metabolism, cholesterol transport; K: Cell adhesion; G and L: n/a. Example proteins were selected based on the statistical significance of the overall difference between protein levels in COVID-19 convalescents and healthy controls (with the adjusted p-value < 0.20 or < 0.05) and based on their relevance for the observed functional enrichment in each cluster. Panel **A**. shows the color-coded protein levels with samples sorted according to race and the presence of Symptoms. Panel B. shows the significance values for two different models. Significance values (adjusted p-values) were transformed as follows: if the observed log_10_-transformed fold change was positive, we calculated 1-p; if negative, we calculated -(1-p). Dark colors indicate adjusted p-value <0.05; light colors adjusted p-value < 0.20; gray: no significance. Columns represent comparisons with significant differences: the Overall difference between COVID-19 convalescents and healthy controls; and the impact of Days since diagnosis, presence of Symptoms, Sex, Age, and Race in a multivariate model using the healthy controls (beige) or COVID-19 convalescents (brown), respectively. The demographic data, protein levels, and results of all models are provided in **Supplementary Data File S1**. D.s.d. - Days since diagnosis; Titer - log_10-_transformed antibody titer as defined in methods; * Less reliable quantitation as determined by a high Coefficient of Variance across quality control samples (>50%)

The five clusters in Figure 4 were enriched in cell adhesion and platelet degranulation (cluster A), innate immunity and the complement system (clusters B and C), hemoglobin, adaptive immunity, and immunoglobulins (cluster D), and adaptive immunity, immunoglobulins, and the complement system (cluster E). Proteins in these five clusters had overall differences between COVID-19 convalescents and healthy controls, but no significant interactions with any known demographics, i.e. no obvious additional biases. The samples were ordered by their annotation as healthy controls or COVID-19 convalescents, as well as by sex and age.

Clusters A to C contain proteins with lower levels in COVID-19 convalescents, e.g. proteins functioning in cell adhesion and platelet degranulation (e.g. fibrinogen A (FGA), VWF), and innate immunity/the complement system (e.g. C1R, C1S, C1QA, C1QC, and GGH)(Figure 4). Actin cytoskeleton proteins (FLNA, PFN1, CFL1, ACTB, TAGLN2, and TSMB4X) showed the strongest bias both with respect to fold-change and significance, as also indicated in Figure 3B. The quantitation of TAGLN2 and TSMB4X quantitation was less reliable as marked in the figure (**Supplementary Data File S1**). In comparison, clusters D and E had proteins with higher levels in COVID-19 convalescents, e.g. immunoglobulins playing a role in antigen recognition (e.g. IGKV1-27, IGKV1-39, IGHV1-46, IGLV3-1, and PRDX2), hemoglobin subunits (e.g. HBB, HBA1, and HBD), and carbonic anhydrase (CA1).

Next, we examined seven clusters with differences between COVID-19 convalescents and healthy controls that were more complex, i.e. that involved interactions with demographic factors (Figure 5). The seven clusters correspond to those also shown in Figure 2 and had the following function enrichments: adaptive immunity, immunoglobulins (clusters F and H), cholesterol transport, lipid metabolism (clusters I and J), cell adhesion (cluster K), and no bias (clusters G and L). Examples shown in Figure 5 were chosen from these pathways.

Age, Sex, Days since diagnosis, or Symptoms had no significant bias with respect to their distribution across healthy controls and COVID-19 convalescents, with the exception of one protein (HPX)(adjusted p-value > 0.20, Figure 5). While this outcome may be in part due to the limited sample size, it might also be attributable to an intrinsic relationships between demographic variables as discussed above (Figure 1D): individuals with symptoms and higher antibody titers during the acute phase tended to have serum samples collected at later time points than those without symptoms and lower antibody titers. We assume that Days since diagnosis and Symptoms/Titer have opposite effects on protein levels: early sample collection and a more severe acute phase of the disease should have similar effects on the serum. Therefore, the inverse relationship between these demographics across samples effectively eliminated any potential signal.

The only demographic variable with an interaction in several clusters was race (Figure 5). To illustrate the effect of race, samples in Figure 5 were sorted into the healthy controls and COVID-19 convalescents, and within the cohorts sorted according to race. The figure focused on samples from only white or African-American (black) individuals which comprised the majority in this study (n=26).

The first three clusters (F to H) in Figure 5 comprised proteins with significantly higher levels in COVID-19 convalescents than in healthy controls, and the difference was stronger when taking race into account. For example, for IGKV3-20, IGKV1-45, IGHG1, IGHV3-49, and IGLV3-27, the difference only existed for black, but not for white individuals. Most proteins from clusters F to H function were immunoglobulins, i.e. they functioned in adaptive immunity.

We observed the opposite race effect in proteins from clusters I to L: proteins had lower levels in COVID-19 convalescents, but more so or only in samples from black individuals (Figure 5). Healthy individuals had no significant biases with respect to race. Proteins from these clusters included those from lipid transport (cluster J), e.g. apolipoproteins (APOB, APOC2, APOC3, APOC4, and APOA4), and cell maintenance proteins, e.g. selenoprotein P (SELENOP)(cluster K). Cluster I showed very weak and mixed interactions with race, with signatures very similar to those in clusters D and E (Figure 4), and it contained proteins from cholesterol transport, e.g. CETP and APOA1.

## Discussion

Proteomic alterations in serum and plasma of mild, moderate, severe and critically ill COVID-19 patients have been studied extensively with respect to changes during the acute phase of the disease ^11,25–30^. In comparison, much less is known about changes upon recovery ^16–19^. We present one of the few controlled studies investigating serum proteomic differences between COVID-19 convalescents and age-, sex-, and race-matched healthy controls. A unique property of the cohort is that samples had been collected early in the pandemic, i.e. from unvaccinated individuals who had the first SARS-CoV-2 infection. However, this opportunity limited cohort size as well as control for various factors, e.g. Days since diagnosis. Further, the complexity of the observed patterns and of the relationship to other findings (see below) limits the discussion to select qualitative observations. ^16^

**Table 1** provides an attempt to relate some of the findings in this study to what is known from literature with respect to serum/plasma proteomic changes during the acute phase of COVID-19. However, it should be noted that due to the vast literature on acute COVID-19, many published observations have contradictory findings for individual proteins/pathways. Example proteins in **Table 1** are listed based on their occurrence in literature. Note that some proteins occur in several rows as they function in several pathways. As available, we also attempt to relate our findings to those on long COVID/PASC in the text below.

While 37 of the 334 proteins (11%) showed significant differences in their levels (adjusted p-value < 0.05, Figure 3), most of the observed proteome was similar between COVID-19 convalescents and healthy controls. **Table 1A** shows pathways and examples from proteins and pathways that were similar between COVID-19 and convalescents and healthy controls, but were observed to be altered during the acute phase of the infection. In comparison, **Tables 1B** and **C** shows pathways and examples of the 37 proteins we found still altered amongst the COVID-19 convalescents, but grouped according to their relationship to literature observations in acute COVID-19: **Table 1B** shows pathways consistent with changes during the acute phase, i.e. proteins from hemolysis, the adaptive immune system and inflammation; **Table 1C** shows pathways whose protein levels were inconsistent with these changes, i.e. proteins from the complement cascade, coagulation, the actin cytoskeleton/cell adhesion/platelet degranulation, and metabolism.

For example, we identified markers of acute inflammation (ORM2) and hemolysis (HBA1, HBB, HBD, and CA1) as well as the hemolytic anemia associated protein PRDX2 with significantly elevated levels in the COVID-19 convalescents compared to healthy controls (Figure 3B, Figure 4), consistent with what had been found in acute COVID-19 patients with high IL-6 levels and severely ill patients ^31^(**Table 1B**). These findings indicated that elevated levels of proteins from inflammation and hemolysis were persistent for up to >2 months of recovery. CA1 is known to associate with the IgA-complex in acute COVID-19 patients but not in healthy individuals ^17^, and the elevated levels we observed indicated that this might still be the case.

Perhaps the most interesting changes were those opposite to the ones observed during the acute phase (**Table 1C**). Examples include proteins involved in coagulation (fibrinogen and VWF) whose levels were significantly reduced amongst COVID-19 convalescents (Figure 4). Unfortunately, as VWF is strongly associated with blood type ^32^, but blood type information was unavailable, the interpretation of these findings are speculative. VWF mediates platelet attachment to damaged endothelium and acts as a carrier protein for coagulation factor VIII rendering protection from proteolytic degradation ^33,34^. Its levels are known to be elevated in acute COVID-19 and long COVID/PASC patients signifying platelet activation and adhesion to endothelium ^33,35^ resulting in COVID-19 associated endotheliopathy ^36^. Fibrinogen regulates protective immune functions and clot formation ^37^. During acute COVID-19, it is involved in thrombosis in lungs ^38^, and fibrinogen chains are strongly associated with COVID-19 fatalities ^39^. The significantly decreased levels of VWF and fibrinogens in COVID-19 convalescents (**Table 1C**) contrasts the finding that most components of the coagulation and complement cascade had returned to pre-infection levels (**Table 1A, Supplementary Figure S5**). One possible explanation is the temporary suppression of some pathways after recovery from the acute phase.

One such pathway relates to platelet counts in the blood. Platelets act as cellular immunomodulators interacting with endothelial cells and leukocytes in response to infections, and are therefore crucial during thrombosis and the host immune response ^40^. During viral infections, low platelet counts, interactions with leukocytes, and platelet secretion may lead to protective or injurious immune effects ^41^. Aberrant blot clot formation, such as thrombosis, is a known complication of COVID-19 infection ^42,43^. As fibrinogen and VWF are engaged in platelet degranulation ^44,45^, we hypothesize that the observed decreased levels of proteins from the platelet degranulation pathway in samples from COVID-19 convalescents (Figure 4, cluster A) may be due to low platelet counts resulting from platelet consumption during COVID-19 infection. This hypothesis is supported by studies that suggest a 5% to 42% exhaustion in platelet counts for several months post infection (immune thrombocytopenia)^46–49^ amongst survivors of both severe and non-severe COVID-19 patients ^50–53^. While low platelet counts can occur any time during the acute phase of COVID-19, it has been frequently observed after clinical recovery ^54^, e.g. after three ^55,56^ or five weeks ^57^ after onset of symptoms, consistent with our findings. Dysregulated platelet function has also been observed in patients of long COVID/PASC ^58,59^.

Similarly, we found significantly reduced levels of proteins of the actin cytoskeleton network amongst COVID-19 convalescents, e.g. PFN1 and CFL1 (Figure 4, cluster A), partially contrasting what had been found for the acute phase (**Table 1C**). The actin cytoskeleton is critical in various pathways of the immune system, ranging from hematopoiesis and immune cell development, recruitment, migration, inter- and intra-cellular signaling, as well as response activation ^60^. Many viruses interact with actin and actin-regulating signaling pathways within the host cell ^61,62^ and reprogram the cellular pathways ^63^. Further, PFN1 is an important player in activation of viral transcription and airway hyperresponsiveness ^64,65^, and it is known to be downregulated in non-severe COVID-19 patients ^64^. CFL1 functions in T cell motility, T cell migration to lymphoid tissues, immune reconstitution, and immune control of viremia ^66^ and is dysregulated in HIV-infected patients ^66,67^. Therefore, an additional interpretation of altered CFL1 levels observed in our data relates to possible changes in T cell mobility.

Further, we observed significant decreases of levels for some proteins of the complement cascade amongst the COVID-19 convalescents (Figure 5) contrasting observations from the acute phase ^68,69^(**Table 1C**). The complement system is also known for a complex relationship with long COVID ^70^: complement dysregulation might even be predictive of long COVID ^71^. The complement cascade directly associates with altered blood coagulation in COVID-19 pathology ^20,72,73^, and blood coagulation, which in turn involves platelet activation.

We discussed the possible impact on platelet counts above which might also explain the temporary depletion of some proteins from the complement system.

Finally, by analyzing the impact of known demographics, we found no significant association of past COVID-19 infections with the individuals’ Age, Sex, Days since diagnosis, and Symptoms. The lack of associations might be due to the small size of the cohort available and the heterogeneity amongst available samples. In comparison, we identified several proteins that were associated with Race which included mostly white and black individuals.

Examples included many proteins of the adaptive immune response (Figure 5). Other proteins with a race effect were from cholesterol metabolism and transport: their elevated levels amongst COVID-19 convalescents were observed more strongly amongst white individuals than black individuals (Figure 5). Apolipoproteins (APOA1 and APOB) are key regulators of cholesterol metabolism and transport ^74,75^ and can render protection against severe COVID-19 ^76–78^. Other apolipoproteins such as APOCs are not known to play a critical role in COVID-19, but demonstrated a race effect in our data (Figure 5). Higher levels of proteins from cholesterol metabolism have also been observed amongst COVID-19 convalescents six weeks after diagnosis ^18^. Hospitalized COVID-19 patients have shown altered lipid metabolism even six months after discharge ^19^.

While grouped into a cluster with proteins with race effects (Figure 5, cluster I), CETP, APOA1, and SAA4 showed individually no significant race difference. CETP is linked to reverse cholesterol transport and associated with APOA1 and SAA4 ^79^. The elevated SAA4 levels are consistent with findings from acute COVID-19 cases ^29,74,80–82;^ however, this is not true for CETP and APOA1 (**Table 1C**). APOA1 and APOA isoforms, which are also involved in the immune response and dyslipidemia, have been frequently observed in COVID-19 patients with acute inflammatory conditions ^29,82,83^. The elevated APOA1 levels contrast our observations on other APOA isoforms which returned to levels similar to those in healthy controls (**Table 1C**) suggesting that further work will be necessary to decipher the complex role of cholesterol metabolism and transport.

Another protein with a race effect was SELENOP (Figure 5). SELENOP is expressed in the liver and secreted into plasma ^84,85^. It has protective functions of host immune defense and tissue homeostasis ^84,85^. SELENOP levels directly impact serum selenium levels ^86^, and higher serum selenium levels, in turn, have been associated with increased COVID-19 survival ^29,86^. We observed elevated SELENOP levels only in the white population but not black individuals which suggest complex, possibly race dependent relationships. In general, while increased COVID-19 infection rates and deaths amongst African American, Hispanic, and Asian communities compared to the white population have been reported ^87^, the race-dependent changes that we observed will require further investigation prior to their interpretations.

In sum, our study provides insights into the proteomic landscape present at up to >two months after the infection. While most of the proteome was similar to that found in healthy individuals, we identified several intriguing differences. The interpretation of these differences, e.g. with respect to a possible temporary decrease in platelet counts, will have to be tested in future work through analysis of larger cohorts. Our findings might inspire some of these analyses to be conducted in a targeted fashion.

## Methods

### Sample collection

All methods were performed in accordance with the relevant guidelines and regulations. We obtained samples from a retrospective case-control study of two cohorts with a total of 58 subjects, comprising healthy controls (n=29) and COVID-19 convalescents (n=29). All participants had been recruited at the University of Georgia at Athens and provided written informed consent prior to participation. The study protocol had been reviewed and approved by the University of Georgia Ethical Review Board (IRB #20202906). The participants’ demographics are shown in **Supplementary Data File S1**. Antibody titers are the maximum dilutions at which antibodies were still detected. For visualization, titers were log transformed (base 10). Serum samples were heat inactivated at 56◦C for 30 min and stored at −80◦C. Samples from COVID-19 convalescents were collected between March-April 2020 based on ‘convenience sampling’ due to the inherent difficulty in access to sample at specific timing at this early point in the pandemic. Therefore, samples were collected without inclusion/exclusion criteria. To construct a retrospective case-control study, we carefully selected healthy control samples based on matching age, sex and race. The healthy controls were selected from an independent cohort at the University of Georgia, Athens (IRB #3773) with demographic information as provided in **Supplementary Data File S1**.

### Sample preparation

Serum samples including individual (n=58) and pooled samples were processed using a protocol described elsewhere ^88^. In brief, 1 μl of serum sample (∼70-80 μg protein) was lysed with 0.1% Rapigest (Waters, MA, USA) in 100 mM ammonium bicarbonate (Sigma, MO, USA) and denatured at 95°C for 5 minutes. Further, the samples were reduced using 5 mM dithiothreitol (DTT, Sigma) at 60°C for 30 minutes, followed by alkylation with 15 mM iodoacetamide (Sigma) at room temperature in the dark for 30 minutes.

Subsequently, the samples were quenched with 10 mM DTT and digested overnight at 37°C with Trypsin gold (Promega, WI, USA). The digestion was stopped and the surfactant was cleaved by treating samples with 200 mM HCl (Sigma) at 37°C for 30 minutes. The samples were desalted on Hypersep C-18 spin tips (Thermo Fisher Scientific, MA, USA) and the peptides dried under vacuum at low heat (Eppendorf, CT, USA). The dried peptides were resuspended in 5% acetonitrile in 0.1% formic acid (Thermo Scientific) and quantified by fluorometric peptide assay kit (Thermo Fisher Scientific) prior to mass spectrometry analysis.

We analyzed the samples using an EASY-nLC 1200 (Thermo Fisher Scientific) connected to Q Exactive HF mass spectrometer (Thermo Fisher Scientific). We used an analytical column RSLC PepMan C-18 (Thermo Fisher Scientific, 2uM, 100Å, 75μm id x 50cm) at 55◦C with the mobile phase comprising buffer A (0.1% formic acid in water) and buffer B (90% acetonitrile in 0.1% formic acid), injecting approximately 400 ng peptides. The chromatographic gradient consisted of 155 minutes from buffer A to buffer B at a flow rate of 300 nl/min with the following steps: 2 to 5% buffer B for 5 minutes, 5 to 25% buffer B for 110 minutes, 25 to 40% buffer B for 25 minutes, 40 to 80% buffer B for 5 minutes, and 80 to 95% buffer B for 5 minutes and hold for additional 5 minutes at 95% for Buffer B.

The serum samples were analyzed using the data independent acquisition (DIA) mode with the following parameters: for full-scan MS acquisition in the Orbitrap, the resolution was set to 120,000, with scan range of 350 to 1650 m/z, the maximum injection time of 100 ms, and automatic gain control (AGC) target of 3e6. The data was acquired using 17 DIA variable windows in the Orbitrap with a resolution set at 60,000, AGC target of 1e6, and the maximum injection time in auto mode.

The order of sample runs was randomized, but we analyzed the age-sex-race matched pairs of healthy controls and COVID-19 convalescents in succession, with a quality control (QC) sample run approximately every 6 samples (**Supplementary Data File S1**). The QC sample consisted of pooled serum samples that had been processed in a way identical to that of the experimental samples.

### Data analysis

#### Primary processing

We used Spectronaut for all primary processing (v14, https://biognosys.com/software/spectronaut/), i.e. identification of fragments from raw mass spectrometry data. All 74 raw files were first converted to the HTRMS format with the HTRMS converter (centroid method). The converted files were then analyzed with directDIA (within Spectronaut) using default settings. We exported intensity information at the fragment level for further preprocessing.

We used in-house R scripts to eliminate the effects arising from samples run at different times. Specifically, sample queuing created four sets, i.e. four batches, and each batch consists of individual serum samples from paired COVID-19 convalescents and healthy controls as well as the QC samples. We first removed fragment ions with values missing in more than half of all samples. We then log_2_-transformed intensity values of the ions and applied Gaussian kernel smoothing with a window of 5 samples: within each batch, we subtracted the time trend captured by Gaussian kernel regression of a kernel width (standard deviation) equivalent to 5 samples. Then, we equalized the median across four batches and the data was transformed back to linear scale. We further applied mapDIA as per published protocol ^89^ to select the best quality fragment ions to estimate the protein levels, and to apply batch-to-batch and within-batch drift normalization to the data at the fragment ion level, prior to deriving values for protein levels. This procedure was part of the mapDIA pipeline. The protein levels were log10-transformed prior to statistical testing as described below.

QC samples were used to assess the variability of protein levels of the 334 proteins, computed as the Coefficient of Variance within each batch (**Supplementary Data File S1**). CoV distributions and principal components separating COVID-19 convalescents and healthy controls, but not the QC samples are shown in **Supplementary Figure S1**. Three QC samples from batch 3 clustered separately from other QC samples. However, samples from COVID-19 convalescents and healthy controls from the same batch clustered correctly with the respective groups suggesting that sample data is accurate. At the raw data level (fragment intensities), linear correlations between QC samples were all >0.968. Proteins with less reliable quantitation are marked in figures with a *.

**Processing for visualization.** For data visualization in heatmaps, the data was standardized by subtracting the row median intensity (across samples) from the intensity value of each protein. The heatmaps were generated using R-scripts and Perseus (version 1.5.5.1)^90,91^. Hierarchical clustering was performed using Perseus setting the complete linkage method and‘1 - Pearson correlation’ as the distance metric. The rows (protein expression signatures) were split into 20 clusters based on similarity with respect to Pearson correlation (Figure 2). Cluster number was chosen based on the total number of proteins in the set. Each cluster was analyzed for enriched functions (NCBI DAVID tool) and example proteins in **Figures 4** and **5** were chosen based on their association with the respective function.

### Statistical testing

#### Overall comparison

To assess the overall difference between protein levels from COVID-19 convalescents and healthy controls, we used a two-tailed, paired t-test. P-values were adjusted for multiple testing correction using the Benjamini-Hochberg procedure ^92^.

#### Linear regression models

To examine the effect of the demographic variables including age, sex, and race, we used three univariate and three multivariate linear regression models. Univariate models considered each variable separately; multivariate models considered each variable in the context of all other variables. All models were evaluated with respect to the variable’s impact on predicting the levels of a specific protein amongst the samples from i) the healthy control data, ii) the COVID-19 convalescents, iii) the data set of paired [log_10_(COVID-19/healthy control)] values. Note that due to the age/sex/race matching, dataset iii) intrinsically controlled for some of the effects of age, sex, and race already. Further, we also considered for healthy controls (dataset i)) variables including Presence of symptoms and Days since diagnosis which were derived from the corresponding sample of the COVID-19 convalescents. We included the variables for control purposes: as their role in healthy individuals is meaningless, we expected no significant associations between the Presence of symptoms and Days since diagnosis when modeling healthy controls (dataset i)). Indeed, we found only a few and minor associations in the univariate models. These associations were likely due to additional links, such as between age and Presence of symptoms. We evaluated the following models: i) Univariate model: protein level (COVID) as a function of Age / Sex / Race / Presence of symptoms / Days since diagnosis; ii) Univariate model: protein level (CONTROL) as a function of Age / Sex / Race / Presence of symptoms / Days since diagnosis; iii) Univariate model: log_10_[protein level ratio (COVID/CONTROL)] as a function of Age / Sex / Race / Presence of symptoms / Days since diagnosis; iv) Multivariate model: protein level (COVID) as a function of Age + Sex + Race + Presence of symptoms + Days since diagnosis; v) Multivariate model: protein level (CONTROL) as a function of Age + Sex + Race + Presence of symptoms + Days since diagnosis; and vi) Multivariate model: log_10_[protein level ratio (COVID/CONTROL)] as a function of Age + Sex + Race + Presence of symptoms + Days since diagnosis + Age*Sex. Due to the correlation between Titer and Symptoms as well as Days since diagnosis, we excluded Titer from the modeling to avoid overfitting. The main text focuses on the results of the multivariate models on CONTROL and COVID sets; the results from all models are presented in the **Supplementary Data File S1**, including corrections for multiple hypothesis testing (see below).

#### Correction for multiple hypothesis testing

We corrected all P-values obtained from the overall comparison and the linear regression models for multiple testing using the Benjamini-Hochberg procedure ^92^. To parse the results, we focussed on proteins with adjusted p-values < 0.05 as the primary set of proteins discussed. We considered an extended set with adjusted p-values < 0.20. All significance values are displayed in gray if not within these thresholds and in dark or light color if below the 0.05 or 0.20 threshold, respectively. Blue and red indicate the directionality of protein level difference. For visualization purposes only, we transformed adjusted p-values (p) to derive a new value as (1-p) if the respective log_10_-transformed fold change was positive, and as -(1-p) if the fold change was negative.

#### Function enrichment and network construction

Using a ranked list of proteins based on directional adjusted p-values of the overall comparison of COVID-19 convalescents vs. healthy controls, we tested for enrichment of protein functions and pathways using Gene Set Enrichment Analysis (GSEA) with the GSEAPreranked tool ^93,94^. To do so, we selected the Human MSigDB’s ^93,95,96^ and the C2 Reactome ^97,98^ databases for function annotations and required 1,000 permutations for analysis. We excluded from the output enriched pathways with >300 or <3 members.

We then used EnrichmentMap v3.3.6 ^99^ within the Cytoscape 3.10.1 package ^100^ to generate an enrichment map displaying the results of the GSEA. Nodes represented significantly enriched functions (false discovery rate < 0.01). Node size in Figure 3 is proportional to the number of genes with the respective annotation. We drew edges between nodes if the two nodes shared a substantial number of genes, i.e. the Jaccard index was ≥0.80. Edge width in Figure 3 is proportional to the Jaccard index.

#### Post-translational modifications

To examine samples for changes in post-translational modifications, we constructed phosphorylation, mono- and di-methylation, a monohexose (+C6+H10+O5) and dihexose (+C12+H20+O10) library with Spectronaut Pulsar (v14, https://biognosys.com/software/spectronaut/), using default settings except for the maximum number of variable modifications set to 3. Then we matched our DIA samples against this library, setting the minor grouping to “by modified sequence” and the differential levels grouping to “minor” (peptide level). We exported peptide intensities from Spectronaut for further analysis. The data is available in **Supplementary Data File S2**.

We extracted peptides with monohexose modifications as well as the corresponding unmodified peptides, as monohexose modifications were the only modifications with substantial numbers of modified peptides. We removed peptides with >20% missing values across the samples. We defined the modification level as log_2_(Intensity_modified_/(Intensity_unmodified_+Intensity_modified_)) for each peptide. We calculated the log_2_ fold change between COVID-19 convalescents and healthy controls and used a paired t-test to determine the significance of the difference. As before, we used the Benjamin-Hochberg procedure to adjust for multiple testing correction.

## Data availability

We deposited the raw files of the serum proteome in the PRIDE database ^101^ with the accession number PXD036597.

## Acknowledgments

CV and TR acknowledge funding by the US National Institutes of Health (75N93019C00052/AI/NIAID). CV acknowledges funding by the US National Institutes of Health (R35GM127089). HC was supported in part by National Medical Research Council, Singapore (NMRC/CG21APR1008). Cohort data were obtained from a study supported by the National Center for Advancing Translational Sciences of the National Institutes of Health under Award Number UL1TR002378.

## Author contributions

Christine Vogel conceptualized this project. Ted M. Ross and Michael A. Carlock provided samples. Shuvadeep Maity, Matthew Pressler, Justin Rendleman, and Burcu Vitrinel analyzed the samples. Smruti Pushalkar, Shaohuan Wu, Hyungwon Choi, and Christine Vogel performed the data analysis and wrote the manuscript. All authors read and approved of the submission of the manuscript.

## Supplementary materials

Supplementary Data File S1.

The file contains demographic data, information on protein level measurements and analysis, as well as order of the mass spectrometry experiments.

Supplementary Data File S2.

The file contains information on peptide level measurements, including post-translational modifications, and analysis of hexose modified peptides.

---

**Figure S1.**
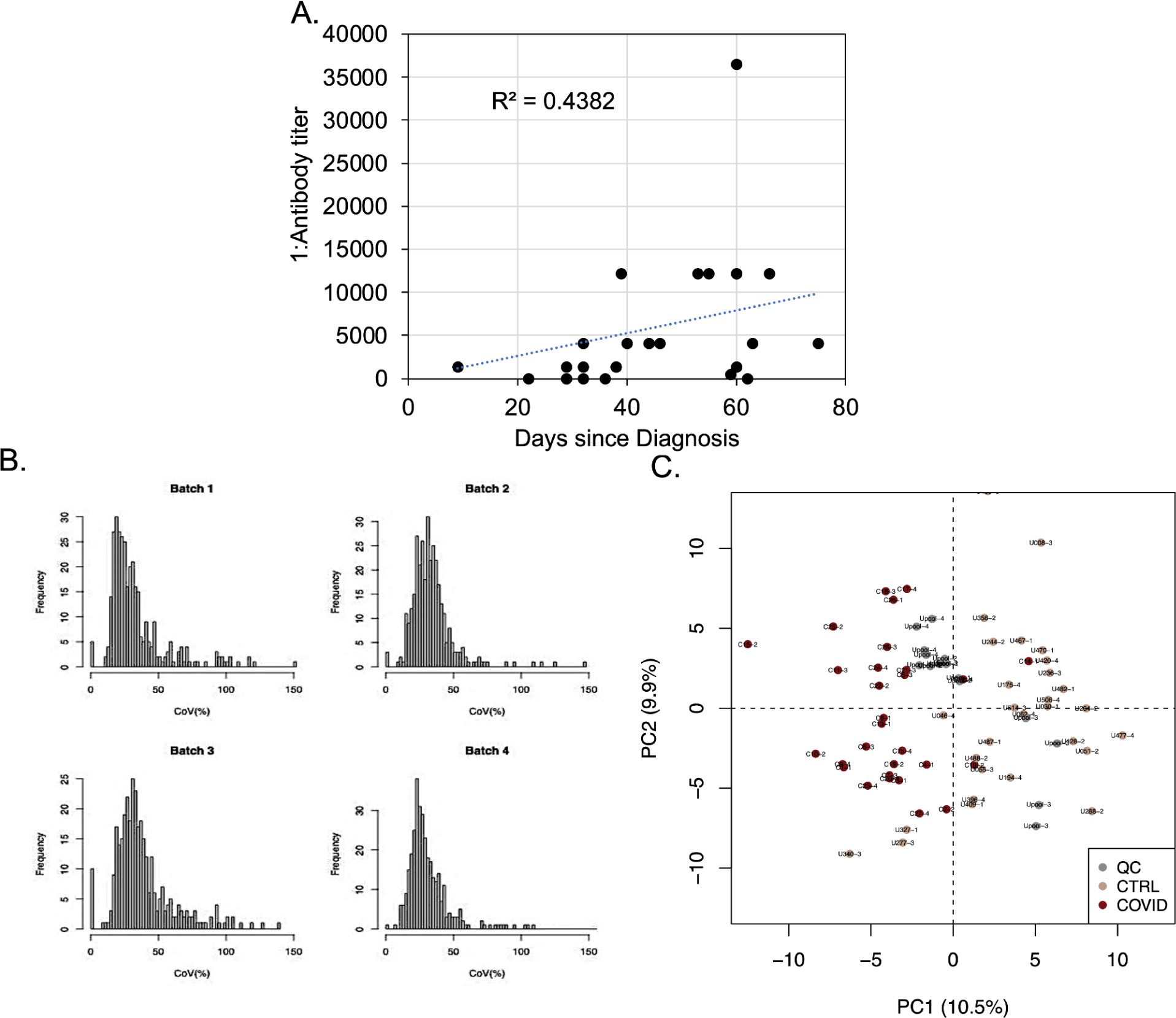
Quality control: fragment intensity variation for quality control samples. We analyzed pooled quality control (QC) samples along with cohort examples (every 6-10 samples). Data was acquired in four sets (batches) listed in **Supplementary Data File S1.** To assess variability of the data, we examined the 334 proteins we discuss in the main text for their variability across the QC samples (labeled ‘Upool’ in the figure). All data was normalized within and across batches (sets) as described in the **Methods**. **A.** Antibody titer levels amongst COVID-19 convalescents (Titer as 1:X) and Days since Diagnosis show substantial correlation. **B.** The four histograms show the distribution of the Coefficients of Variance (CoV) for the protein levels grouped into the four technical batches. Almost all proteins (330 of 334) have <50% CoV in at least one set which is in the expected range for untargeted (shotgun) proteomics. We did not filter for CoV but marked the few proteins of CoV>50% in Figure 3B and in the main text. **C.** The plot shows all samples, including the QC samples (‘Upool’), after normalization in their distribution across the first two principal components. This figure is analogous to **Figure 3A**. The Healthy controls and COVID-19 convalescents are clearly separated and QC samples cluster in the center. A minor exception are QC samples from technical batch 3 which group locate off center; however COVID-19 convalescents and Healthy control samples from batch 3 cluster correctly with their respective groups, indicating successful data acquisition and normalization.

**Figure S2.**
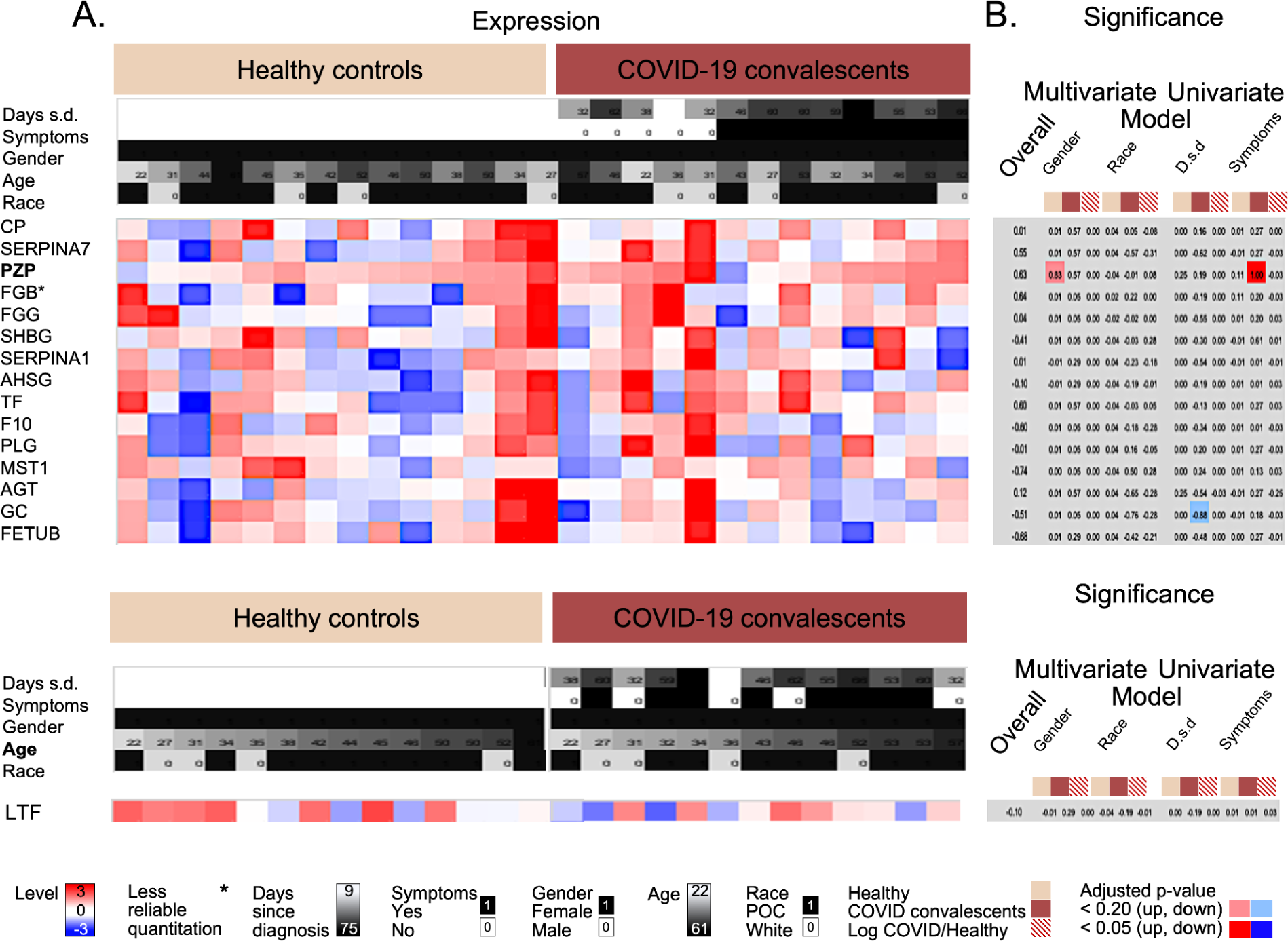
Quality control: expected expression levels of female-specific proteins. Heatmap showing levels of Pregnancy Zone Protein (PZP) along with proteins with similar patterns as well as Lactotransferrin (LTF). **A.** The panel shows the color-coded protein levels for female individuals from both cohorts. **B**. The panel shows the significance values for different comparisons and different models. Significance values (adjusted p-values) were transformed as follows: if the observed log_10_-transformed level fold change was positive, we calculated 1-[adjusted p-value]; if negative, we calculated -(1-[adjusted p-value]). Dark colors indicate adjusted p-value <0.05; light colors adjusted p-value < 0.20. Columns represent select comparisons: the Overall difference between COVID-19 convalescents and healthy controls; the role of sex and race in a multivariate model in which other factors such as age were also considered; and the role of Days since diagnosis and the presence of Symptoms in a univariate model which considered only one factor at a time. Each statistical model was developed separately for the factor in question and the respective sample set: healthy controls (beige), COVID-19 convalescents (brown); log_10_-transformed ratio of paired protein levels for COVID-19 cases and healthy control (beige-brown striped). The complete results of the statistical testing are provided in **Supplementary Data File S1**. Importantly, PZP is the only protein of the 334 total proteins with a sex difference: it has slightly higher levels in healthy women than inhealthy men (adjusted p-value < 0.20). As PZP levels are thought to associate with pregnancy, the results suggest that some of the female individuals were pregnant, causing the difference in levels between the two sexes. Further, while not significant, LTF shows a general decline in levels with age of healthy female individuals, concurrent with the role of LTF in breastmilk. POC - person of color

**Figure S3.**
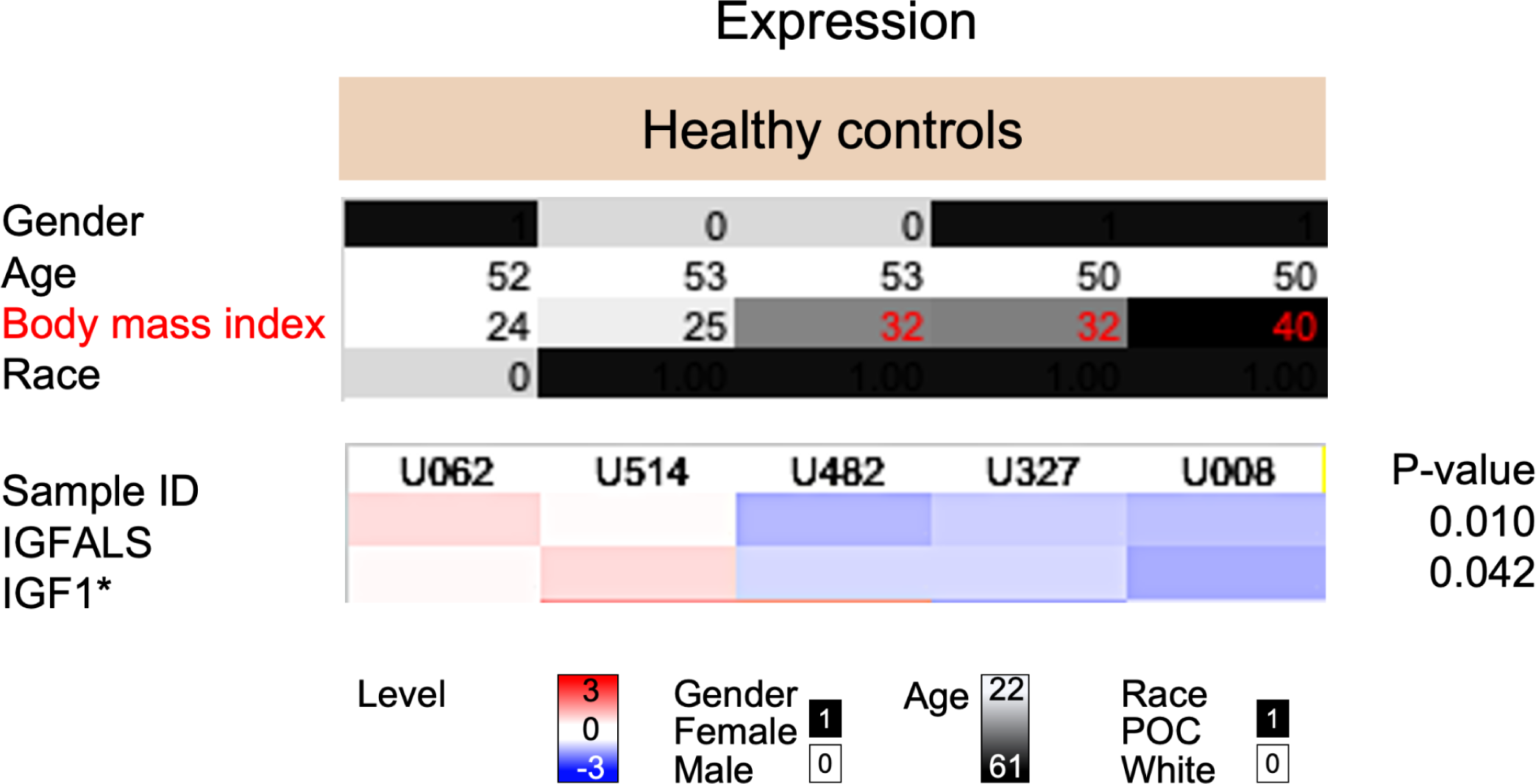
Quality control: expected expression patterns of obesity related proteins. The figure shows the meta data and protein levels for IGFALS and IGF1for 5 healthy control individuals. Both proteins have been connected to obesity. The 5 individuals were selected based on similar age (50 to 53 years) but different body mass index (BMI): normal weight (BMI<25) versus obese (BMI>30). Due to the small size of the cohort, there were no additional sets of age-matched individuals from different BMI categories. We found a significant difference in IGFALS and IGF1 levels between normal weight and obese individuals (P-value, 0.05, two-sample t-test). The demographic information and protein level data are provided in Supplementary Data File S1. POC - person of color

**Figure S4.**
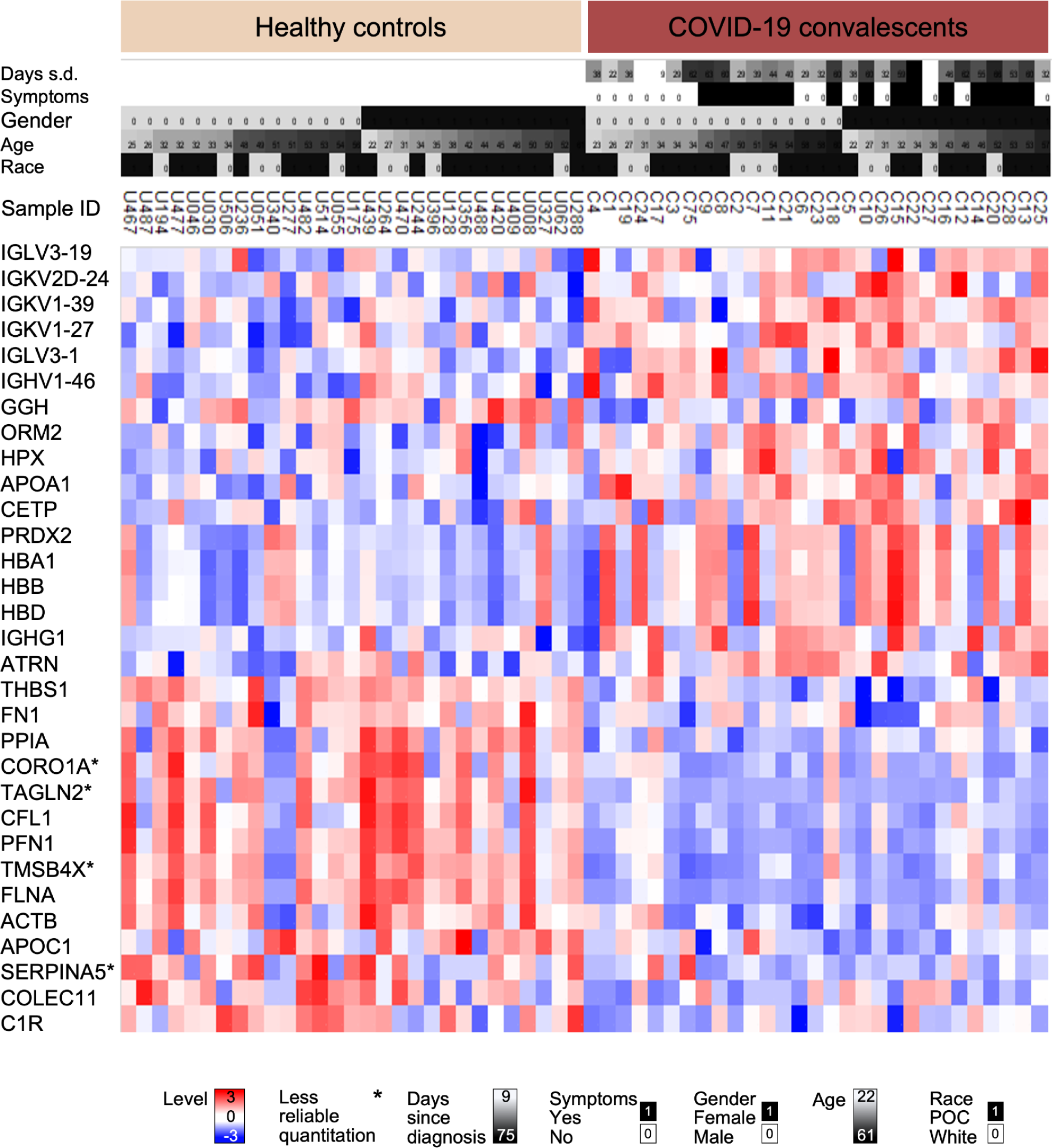
Proteins with statistically different levels between COVID-19 convalescents and healthy controls. Shown are the normalized protein levels for the statistically significantly expressed proteins (adjusted p-value < 0.05). All data is available in **Supplementary Data Files S1**. Samples are organized according to sex and age. Proteins with high Coefficient of Variance are marked with * (>50%). POC - person of color; S.d. - since diagnosis.

**Figure S5.**
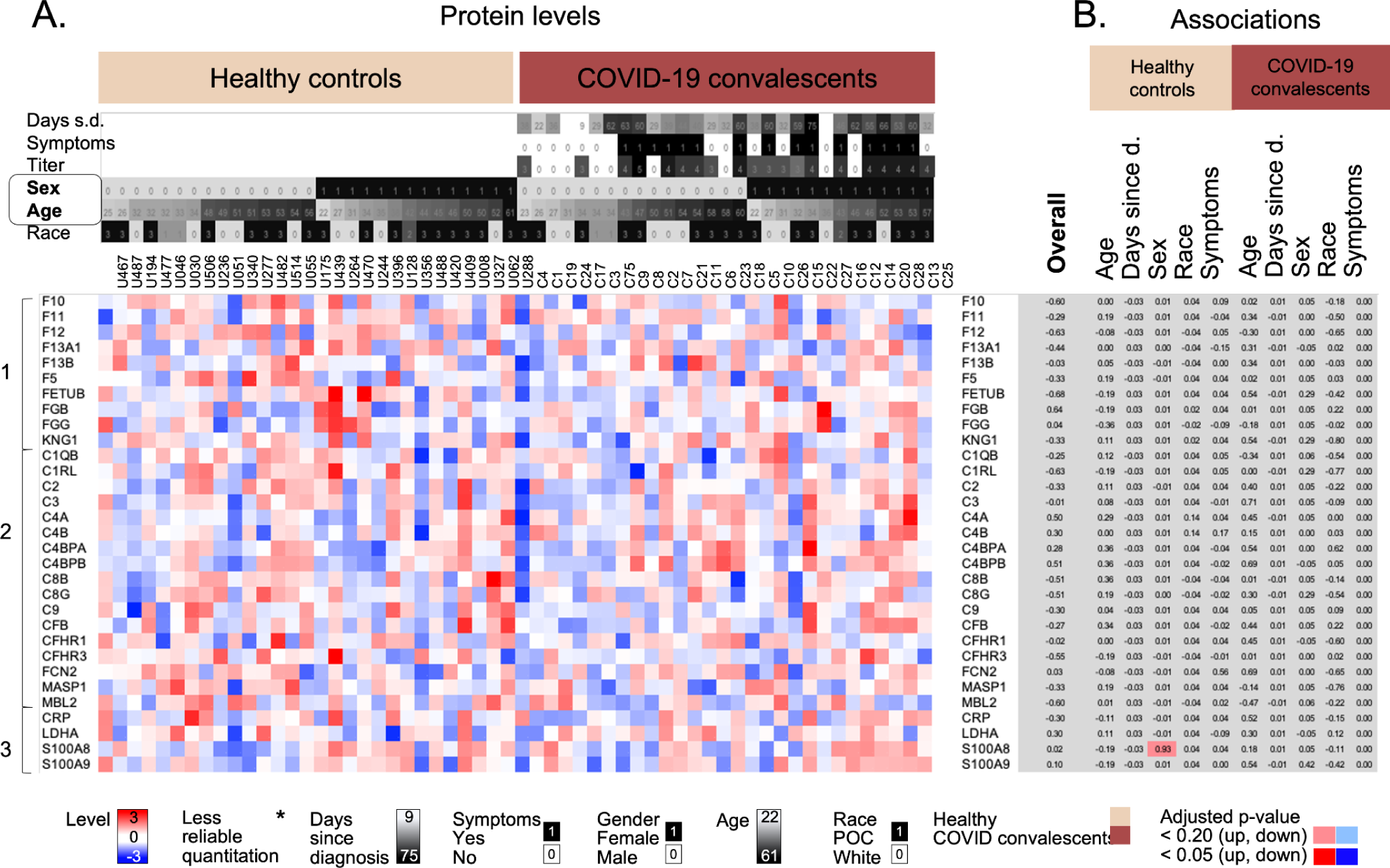
Proteins whose levels were similar between COVID-19 convalescents and healthy controls. Heatmap showing proteins from three pathways that are largely similar in healthy controls and COVID-19 convalescents. **A.** The panel shows the color-coded protein levels with samples sorted according to sex and age. **B.** The panel shows the significance values (associations) for different comparisons and different models. Significance values (adjusted p-values) were transformed as follows: if the observed log_10_-transformed fold change was positive, we calculated 1-p; if negative, we calculated -(1-p). Dark colors indicate adjusted p-value <0.05; light colors adjusted p-value < 0.20; grey: no significance. Columns represent select comparisons: the ‘Overall’ difference between COVID-19 convalescents and healthy controls; and the impact of Days since diagnosis, the presence of Symptoms, Sex, Age, and Race in a multivariate model using the healthy control (beige) or COVID-19 convalescents (brown). The complete results of the statistical testing are provided in **Supplementary Data File S1**. Clusters: 1 Coagulation cascade; 2 Complement system; 3 Inflammation.

**Figure S6.**
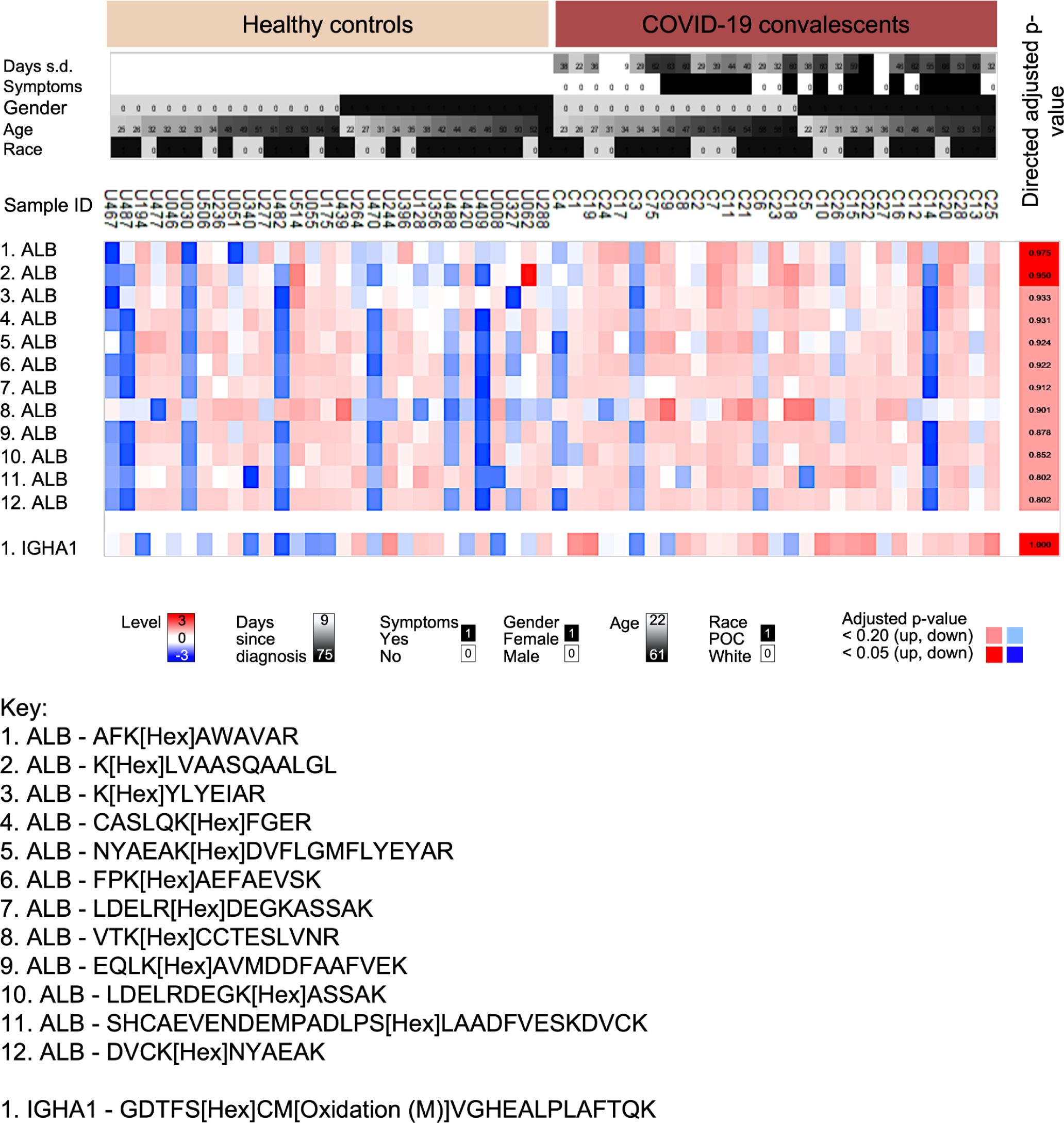
Post-translational modifications. The heatmap shows examples of normalized levels of hexose modified peptides. Three peptides (two for albumin (ALB) and one for Immunoglobulin heavy constant alpha 1 (IGHA1) were significantly more modified amongst COVID-19 convalescents compared to healthy controls (adjusted p-value < 0.05). The heatmap shows additional modified peptides with an adjusted p-value < 0.20. For visualization, data was row median centered and row Z-score normalized. While highly heterogeneous, healthy controls show more often low levels of modification than COVID-19 convalescents, in particular amongst women. The peptide sequence is shown in the key below the heatmap.

**Table S1:**
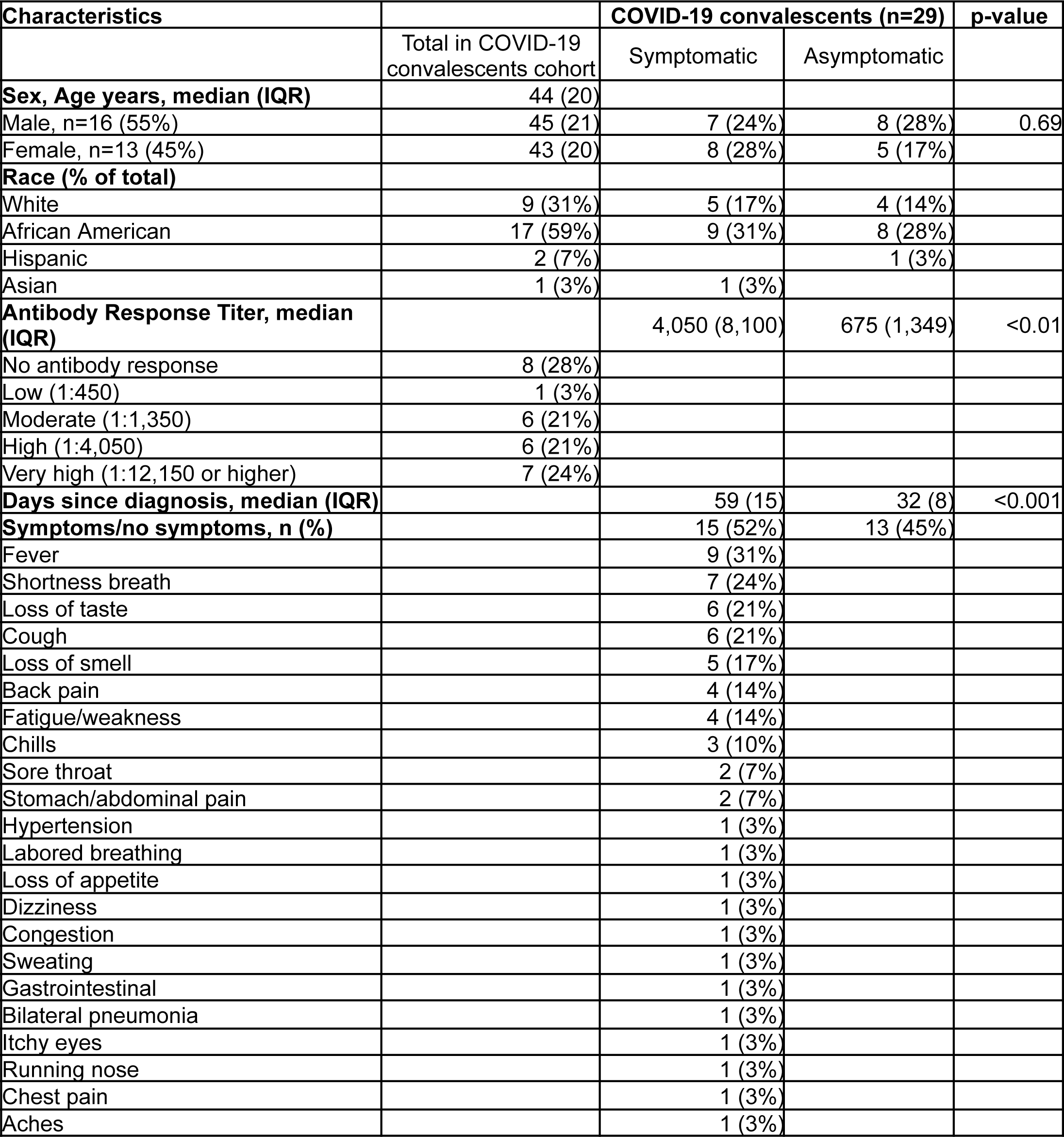
Demographics of COVID-19 convalescents. Demographic data per healthy and convalescent subject are provided in **Supplementary Data File S1.** IQR - interquartile range.

